# Mitomycin C Retains Efficacy after Adaptive Laboratory Evolution of *Staphylococcus aureus*

**DOI:** 10.1101/2024.10.14.618284

**Authors:** Maiken Engelbrecht Petersen, Amanda Batoul Khamas, Lars Jørgen Østergaard, Nis Pedersen Jørgensen, Rikke Louise Meyer

## Abstract

Antibiotic resistance is one of the greatest threats against human health and the misuse and overuse of antibiotics is a key factor driving resistance development. During prolonged antibiotic treatment of chronic infections, the antimicrobial pressure facilitates selection of antibiotic resistance mutations. It has been suggested that using antibiotics in combinations may reduce the emergence of resistance. Furthermore, antibiotic tolerant persister cells may be a reservoir for resistance development, so targeting persister cells with anti-persister drugs could also reduce the emergence of resistance.

In this study, we conducted a 42-day adaptive laboratory evolution experiment using *Staphylococcus aureus* exposed to common antibiotics and the anti-persister drug mitomycin C, either alone or in combination. We monitored susceptibility daily and assessed phenotypic changes in growth and biofilm formation in evolved strains. Whole-genome sequencing revealed mutations linked to antibiotic resistance and phenotypic shifts.

Resistance developed rapidly against rifampicin, while ciprofloxacin and daptomycin showed slower resistance emergence. Treatments with vancomycin or mitomycin C resulted in minimal changes in susceptibility. Combination therapies generally delayed resistance, though resistance was not fully prevented. Notably, mitomycin C combined with rifampicin effectively suppressed rifampicin resistance. Sub-inhibitory antibiotic concentrations were associated with both known and novel mutations, including in the nucleotide excision repair system and azoreductase, following mitomycin C treatment—mutations not previously reported.

While combination therapy delayed resistance, mitomycin C’s efficacy and ability to prevent rifampicin resistance highlights its potential in combating antibiotic resistance. Further investigation is needed to evaluate the broader application of anti-persister drugs in resistance prevention.

## Introduction

Antibiotic resistance has been named one of the greatest threats against human health in the coming decades (1), and it is already the second most prevalent cause of death from infections (2). Resistance emerges when bacteria are exposed to antibiotics in various settings, such as hospitals, in livestock production, and in environments contaminated with pharmaceutical wastewater (3). Some of the most prevalent antibiotic-resistant bacteria in hospital-acquired infections are methicillin resistant *Staphylococcus aureus* (MRSA) and vancomycin intermediate *S. aureus* (VISA) (4, 5). The first-line treatment against MRSA infections is vancomycin or daptomycin therapy, and combination therapy with rifampicin is frequently administered if the infection is associated with an implant (6). Resistance to any of these drugs significantly complicates the treatment. Although daptomycin resistance is rare, it has been reported (7). Intermediate vancomycin resistance is more common (8), and a single study has also reported a daptomycin resistant, vancomycin intermediate MRSA isolate (9).

Development of antibiotic resistance is often associated with long-term antibiotic treatment of recalcitrant infections, such as implant-associated infections that involve biofilms. To increase the antimicrobial potency, antibiotic combination therapy is often used to treat this kind of infection (6, 10). In theory, combining antibiotics that have different molecular targets also reduces the risk of resistance development, since more mutations are needed to gain resistance (5). For this reason, rifampicin is rarely used as monotherapy because resistance to this drug emerges easily (6). However, in practice, combination therapy does not always protect against the emergence of resistance (11, 12). The success of this strategy most likely depends on the drug’s mode of action, the number of mutations required for resistance to occur, and the ability of the drugs to act in synergy against the pathogen.

One of the most common mechanisms of antibiotic resistance is modification of the drug’s molecular target, resulting in decreased binding affinity (13). Antibiotics that target a protein are most prone to resistance development, as a single point mutation may alter the binding affinity. This is the case for e.g. rifampicin (14). In contrast, other cellular structures, such as the cell membrane (the target of daptomycin) or the cell wall (the target of vancomycin) require more complex structural alterations to convey resistance through target modification. From this perspective, DNA represents a potentially attractive target for antibiotics. However, DNA is obviously not unique to bacteria, and DNA-targeting antimicrobials will therefore also be cytotoxic. Vice versa, DNA-targeting antineoplastic drugs used in cancer therapy also have antimicrobial activity, and recent research has pointed to mitomycin C as a powerful antimicrobial. Mitomycin C is a chemotherapeutic drug, currently approved for treatment of several types of malignant cancers. It has gained increasing attention during the last decade as a potential candidate for drug repurposing to combat recalcitrant bacterial infections (15–18). Unlike most antibiotics, mitomycin C kills bacteria independently on metabolic activity, as it crosslinks DNA leading to cell death (19). This is a favourable trait as it makes mitomycin C effective at eradicating bacterial persister cells (17) – subpopulations of bacteria with high antibiotic tolerance due to their transient inactive state (20). This phenotype is associated with implant-associated infections where bacteria reside in biofilms, out of reach from the immune system (21, 22). Implant-associated infections therefore require months or years of antibiotic treatment if surgical intervention is impossible (23). Due to the lengthy treatment of these infections, persister cells have been linked to the emergence of antibiotic resistance (24).

We hypothesize that using antimicrobials, such as mitomycin C, to kill persister cells will prevent resistant mutants from emerging due to the shorter treatment time. We also hypothesize that mitomycin C is less prone to resistance development because its molecular target (DNA) cannot change its drug-binding affinity through point mutations, and because it has the same antimicrobial activity against all bacterial cells in a population.

The aim of this study was to determine if *S. aureus* can develop resistance against mitomycin C, and if combination therapy with rifampicin prevents or delays resistance against this drug as well as other antibiotics. To address these aims, we performed an adaptive laboratory evolution experiment using four clinically relevant antibiotics (ciprofloxacin, vancomycin, daptomycin, rifampicin) and mitomycin C in monothSerapy or as combination therapy with rifampicin. We then identified which genomic mutations occurred during adaptive laboratory evolution to generate resistance, and how these mutations affected the general phenotype of the evolved strains.

## Results

### Adaptive laboratory evolution significantly decreases antibiotic susceptibility for all tested antibiotics except mitomycin C

We studied the emergence of antibiotic resistance in *S. aureus* exposed to antibiotics in monotherapy and combination therapy in an adaptive laboratory evolution experiment over 42 days, generating three independently evolved strains for each antibiotic treatment. In adaptive evolution, bacteria are inoculated into a range of antibiotic concentrations, and bacteria from the highest antibiotic concentration that allowed growth is used to inoculate the assay each day, selecting for higher and higher resistance to the drug.

During the 42 cycles of adaptive evolution, MIC values increased for all antibiotic monotherapies. The MIC for rifampicin increased quickly resulting in >128.000 fold increase after just 7 days (Figure 1A, Table 1). The increase in MIC for other antibiotics was more gradual, but by the end of the 42 days, MIC values had increased by 4 fold for mitomycin C, 8 fold for vancomycin, up to 128 fold for daptomycin, and up to 1024 fold for ciprofloxacin (Figure 1A, Table 1). However, the MIC values for MMC and vancomycin dropped down after sub-culturing leading to an overall increase in MIC by 2-fold and 4-fold, respectively. The clinical breakpoints for resistance is 0.5 μg/ml for rifampicin, 1 μg/ml for ciprofloxacin, and 2 μg/ml for vancomycin. There are no breakpoint values available for daptomycin and mitomycin C. Breakpoint values are based on several factors: MIC values (measured in Müller Hinton broth) and considerations about pharmacokinetic and pharmacodynamic aspects of the drug (25). The breakpoint/MIC ratios for different pathogens and antibiotics therefore vary from 1-250 among susceptible strains (26). Our aim was to investigate the onset of resistance, and to reflect this, we chose to define resistance as occurring when an evolved strain reached a ≥4-fold increase in MIC compared to the parent strain, no matter what the breakpoint value was for the drug of interest. Using this threshold, resistance developed almost immediately for rifampicin, within a week for ciprofloxacin, after approx. 30 days for vancomycin, after approx. 20 days for daptomycin, and never for mitomycin (Table 2).

**Table 1.** MIC values of evolved strains. MIC post-evolution was determined from the final cycle of the adaptive laboratory evolution. For combination therapies, pre-and post-evolution MICs were measured in the presence of rifampicin and therefore notes the rifampicin also. MIC “post-evolution and culturing” was determined for single antibiotics after removing antibiotic pressure overnight prior to inoculating the MIC assay. The final “fold increase” in MIC was calculated from MIC values of the parent strains in the absence of rifampicin, and MIC values of evolved strains in the absence of rifampicin. Several values are shown when the 3 replicate evolved strains did not display the same MIC. In these cases, the numbers are in respective order of strains 1, 2 and 3. All values are in μg/mL. *The maximum concentration of rifampicin that the samples were exposed to during adaptive laboratory evolution.

|  | Strain evolved with | Antibiotic tested | MIC of parent strain | MIC post-evolution | MIC post-evolution and culturing | Fold increase (monotherapy relative) | Max. rif. exposure conc.* |
| --- | --- | --- | --- | --- | --- | --- | --- |
| Monotherapy | MMC | MMC | 0.4 | 1.6 | 0.8 | 2 | - |
|  | Vanco | Vanco | 2 | 16 | 8 | 4 | - |
|  | Cipro | Cipro | 1 | 128, >1024, >1024 | 128, >1024, 1024 | 128, >1024, 1024 | - |
|  | Dapto | Dapto | 2 | 1024, 1024, 256 | 512, 512, 128 | 256, 256, 64 | - |
|  | Rif | Rif | 0.008 | >1024 | >1024 | >128000 | 1024 |
| Combination therapy | MMC + Rif | MMC<br>Rif | 0.2<br>0.008 | 1.6<br>0.016 | 0.8<br>0.031, 0.063, 0.016 | 2<br>4, 8, 2 | 0.016 |
|  | Vanco + Rif | Vanco<br>Rif | 1<br>0.008 | 8<br>0.063 | 8<br>8, 8, 2 | 8<br>1024, 1024, 256 | 0.063 |
|  | Cipro + Rif | Cipro<br>Rif | 0.5<br>0.008 | 64<br>0.25 | 32<br>4, 4, 2 | 32<br>512, 512, 256 | 0.5,<br>0.25,<br>0.25 |
|  | Dapto + Rif | Dapto<br>Rif | 1<br>0.008 | 256, 16, 32<br>8, 0.25, 1 | 256, 16, 16<br>512, 1, 2 | 128, 8, 8<br>65536, 128, 256 | 8, 0.25,<br>1 |

**Table 2.** Days of evolution before resistance emerged. Resistance was defined as reaching a >4-fold increase in MIC. When only one value is written, the number is identical between the three evolved strains. Three values are shown when results from the three replicates differed.

|  | Antibiotic | Days before<br>developing resistance |
| --- | --- | --- |
| Mono-<br>therapy | Mitomycin C | - |
|  | Vancomycin | 33, 31, 29 |
|  | Ciprofloxacin | 6, 7, 6 |
|  | Daptomycin | 22, 24, 20 |
|  | Rifampicin | 3 |
| Combination<br>therapy | Mitomycin C | - |
|  | Vancomycin | - |
|  | Ciprofloxacin | 18, 22, 23 |
|  | Daptomycin | 27, 27, 31 |

**Figure 1.**
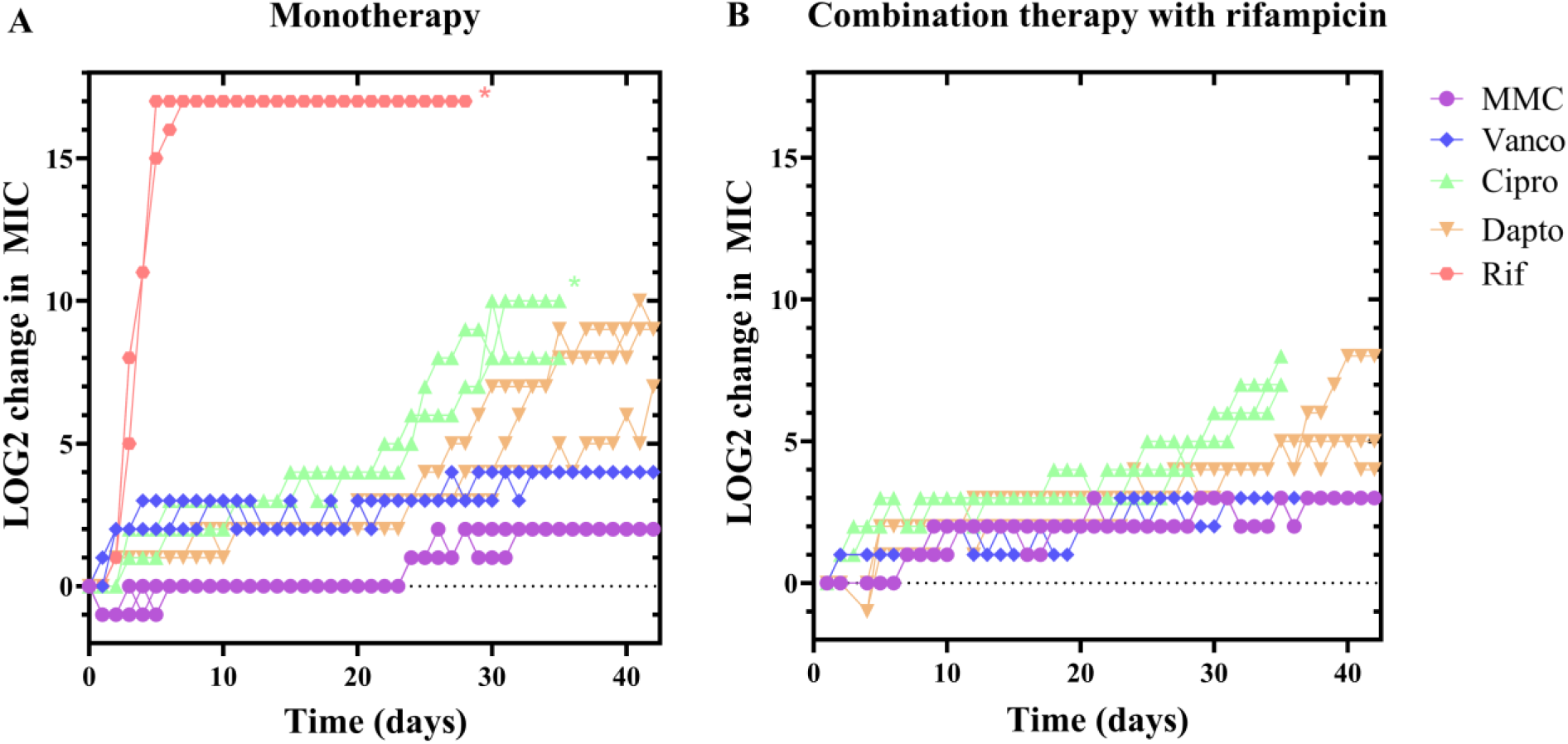
Resistance development during continuous sub-inhibitory antibiotic treatment. *S. aureus* was grown to mid exponential phase and transferred to 96-well plates containing dilution series of either MMC, Vanco, Cipro, Dapto or Rif monotherapy **(A)** or combination therapy **(B)** in combination with rifampicin. The plates were incubated at 37 °C overnight and the MIC was determined by reading plates at 600 nm. Bacteria from wells with antibiotic concentrations immediately below MIC were transferred to a new antibiotic dilution series of the same antibiotic treatment as they had previously received, and incubated at 37 °C overnight. This cycle was repeated 42 times. As the MIC increased, the concentration range of antibiotics was adjusted include concentrations from concentrations from 0.25 to 2 fold the MIC value from the previous cycle. n = 3 independent evolutions for each treatment. *The treatment reached the upper limit of detection.

**Figure 2.**
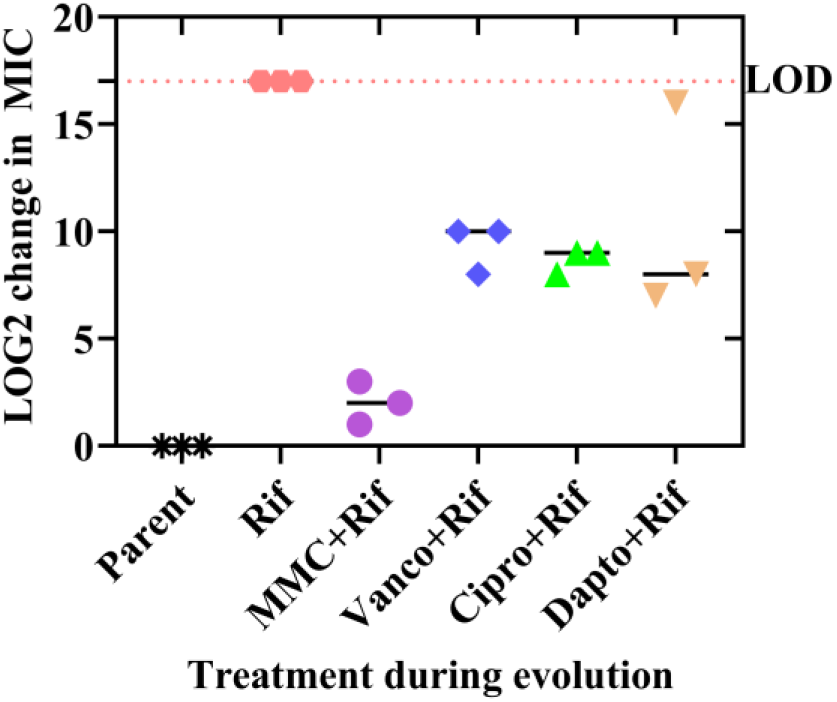
Rifampicin MIC post-evolution of strains receiving combination therapy. Evolved strains were incubated in a rifampicin dilution series in TSB and incubated overnight. The rifampicin MIC was normalised to the MIC of the pre-evolution strain (parent). LOD: Limit of detection, the maximum obtainable rifampicin concentration (1024 μg/mL).

Adaptive evolution was conducted in three independent replicates for each treatment, and resistance developed almost simultaneously in the replicates. The pattern was only heterogenous in cultures exposed to daptomycin, indicating that changes in daptomycin susceptibility involves several steps. Furthermore, MIC values for daptomycin fluctuated from day to day rather than increasing after each cycle, which could indicate transient antibiotic tolerance rather than resistance (27).

In combination therapy with rifampicin, the MIC for other antibiotics still increased during adaptive evolution, but it generally took longer before resulting in resistance (Figure 1B, Table 2). For some antibiotics (ciprofloxacin and daptomycin) MIC values at the end of the experiment were lower than for strains evolved under monotherapy (Table 1). However, they were still much higher than the MIC for the parent strain, and we therefore conclude that combination therapy with rifampicin delayed resistance, rather than preventing it.

### Combination therapy with mitomycin C prevents development of rifampicin resistance and does not lead to cross-resistance

Our primary objective was to determine if rifampicin prevented or delayed development of resistance to other antibiotics. However, we also measured the MIC of rifampicin after 42 days of combination therapy, and found that it was much lower in strains evolved under combination therapy rather than monotherapy (Table 1). It should be noted that the selective pressure of rifampicin was higher in rifampicin monotherapy compared to combination therapies, and this difference may explain the result (see right-most column in Table1). However, it is also possible that combination therapy will delay or attenuate rifampicin resistance. Most impressively, the MIC for rifampicin remained below the clinical breakpoint during combination therapy with mitomycin C, and rifampicin MIC only increased 4-, 8- and 2-fold for the three strains receiving MMC and rifampicin combination treatment, compared with rifampicin monotherapy that rapidly increased MIC >128000-fold (Table 1). The mechanism-of-action of mitomycin C is covalent crosslinking of DNA (19), however, it is not mutagenic (28) and has even been described as inhibiting mutagenesis (29), which may explain the lack of rifampicin resistance development seen here.

During evolution of resistance, it is not uncommon that resistance against one antibiotic causes resistance against another. Such cross resistance has e.g. been observed in *E. coli* after evolution using sub-inhibitory concentrations of the chemotherapeutic drug bleomycin (30). Since this is the first study to investigate evolution of resistance to mitomycin C, we also investigated if evolution under selective pressure from mitomycin C could lead to cross resistance. All three strains evolved under mitomycin C monotherapy had the same MIC for vancomycin, daptomycin, and ciprofloxacin as the parent strains, and the MIC for rifampicin only increased two-fold (Table 3). Therefore, 42 days of exposure to mitomycin C did not lead to mitomycin C resistance or any cross resistance.

**Table 3.**
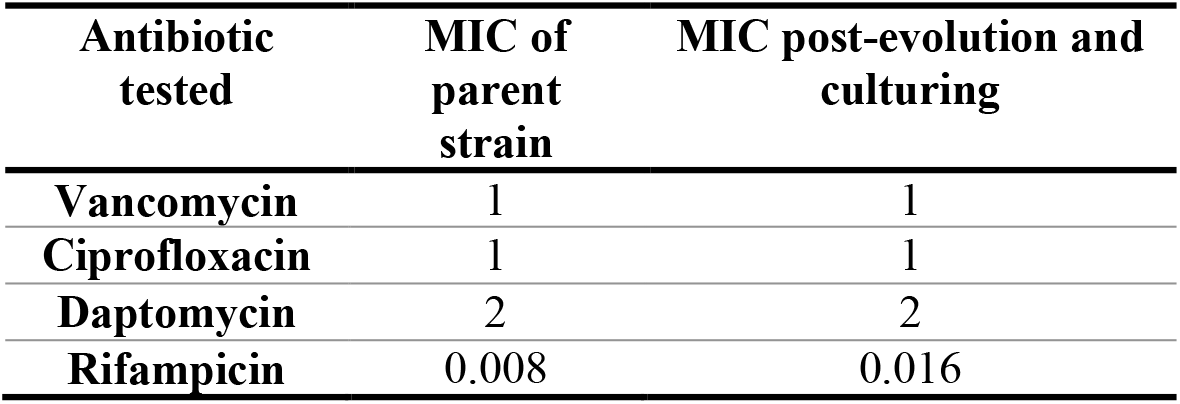
MIC values of strains evolved with mitomycin C. To test for cross resistance, MIC was performed on the three strains receiving mitomycin C during 42 cycles of adaptive evolution and compared with parent strains.

| Antibiotic<br>tested | MIC of<br>parent<br>strain | MIC post-evolution and<br>culturing |
| --- | --- | --- |
| Vancomycin | 1 | 1 |
| Ciprofloxacin | 1 | 1 |
| Daptomycin | 2 | 2 |
| Rifampicin | 0.008 | 0.016 |

### Evolved strains display phenotypic changes in growth pattern and biofilm formation

Development of antimicrobial resistance is sometimes accompanied by changes in growth patterns that partly explain the ability of the evolved strain to survive exposure to antibiotics. We therefore characterized the planktonic growth rates and ability to form biofilm in all of the evolved strains. For all evolved strains, the growth rate either decreased or was unaltered compared to the parent strain (Figure 3A, Table 4). The evolved strains with the slowest growth rates were MMC + Rif strain 2, Dapto strain 1 and 2, and Dapto + Rif strain 1(Table 4). Interestingly, the most slow-growing strains evolved under daptomycin exposure (Dapto 1, Dapto 2 and Dapto + Rif 1) also displayed the highest MICs (512, 512, 128 μg/ml, respectively) relative to the fast-growing strains (Dapto 3, Dapto + Rif 2 and Dapto + Rif 3, with MICs of 128, 8, 8 μg/ml). Some evolved strains displayed a biphasic growth curve. This phenomenon was most pronounced in strains evolved under daptomycin monotherapy, or rifampicin combination therapy with daptomycin or vancomycin. The biphasic growth curve usually indicates a temporary growth arrest after glucose is exhausted from the media (31). Their appearance in evolved strains is indicative of changes in control of metabolic pathways, or in the activation of the stringent response, which has been linked to the sudden growth arrest at the point of glucose exhaustion from a complex media (32).

**Table 4.** Growth rates for evolved strains and parent strain. Two measurements were made for each strain (in parentheses) expect for the parent strain where n = 6. Growth rates were calculated from growth curves (Figure 3a).

|  | Strain | Growth rate (au) |
| --- | --- | --- |
|  | Parent | 1.50 ± 0.03 |
| Monotherapy | Mitomycin C | (1.32, 1.32), (1.40, 1.40), (1.39, 1.39) |
|  | Vancomycin | (1.29, 1.30), (1.24, 1.25), (1.32, 1.32) |
|  | Ciprofloxacin | (1.22, 1.22), (1.39, 1.41), (1.46, 1.48) |
|  | Daptomycin | (1.08, 1.08), (1.18, 1.18), (1.35, 1.36) |
|  | Rifampicin | (1.27, 1.27), (1.45, 1.44), (1.44, 1.45) |
| Combination therapy | Mitomycin C | (1.37, 1.33), (1.19, 1.20), (1.46, 1.48) |
|  | Vancomycin | (1.31, 1.31), (1.37, 1.37), (1.29, 1.29) |
|  | Ciprofloxacin | (1.42, 1.41), (1.22, 1.25), (1.21, 1.23) |
|  | Daptomycin | (1.11, 1.10), (1.35, 1.35), (1.38, 1.38) |

**Figure 3.**
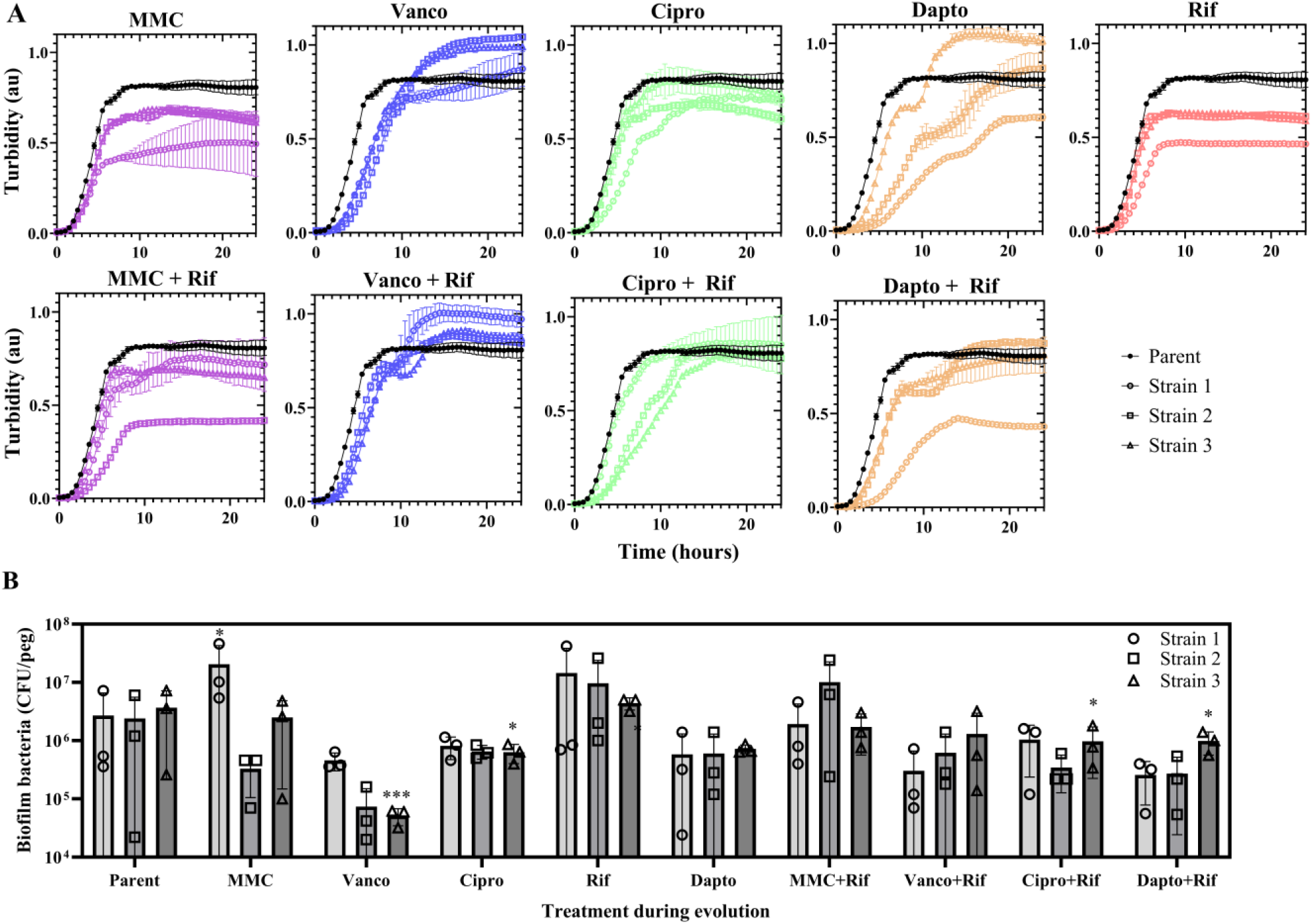
*S. aureus* growth and biofilm formation before and after adaptive evolution. (**A**) *S. aureus* evolved with mitomycin C (MMC), vancomycin (Vanco), ciprofloxacin (Cipro), daptoymycin (Dapto), rifampicin (Rif) monotherapy or in combination therapy with rifampicin were grown in TSB in a 96-well plate at 37°C, and turbidity was measured every 30 min for 24 h. Parent strains are shown in black (mean, n =6, error bars = standard deviation) and evolved strains in colour (n=2). (**B**) Biofilm formed on peg lids in TSB for 3 days media exchange. Viable cells in biofilms were quantified as CFU. Bars show means with standard deviation and symbols show the individual datapoints for each replicate sample (n = 3 for each independently evolved strain).

Next, we investigated the ability of the evolved strains to form biofilm in rich media under static conditions. A few evolved strains produced less biofilm, but there was no trend among strains evolved under the same treatment (Figure 3B). Only a single strain (MMC strain 1) displayed increased biofilm formation.

### Resistance to rifampicin, ciprofloxacin, vancomycin, and daptomycin was linked to the emergence of well-known mutations

After 42 cycles of adaptive evolution, full genome sequencing was performed on the 27 evolved strains and three parent strains to identify mutations that may have caused the resistant phenotype. We assembled the genomes against a reference genome and performed variant calling to identify single nucleotide polymorphisms (SNPs). We compared SNPs identified in the evolved strains with SNPs in the parent strains to identify mutations that had occurred during adaptive evolution. After identifying the genetic mutations, “key mutated genes” were defined as genes that have previously been known to cause resistance, or genes that were mutated at least twice (in the same strain or in two replicate strains) during evolution under a specific antibiotic pressure.

Rifampicin resistance is well characterized and usually associated with mutations in *rpoB* coding for the RNA polymerase β –subunit (14, 33). We therefore expected to find mutations in *rpoB* in all strains that developed rifampicin resistance. Indeed, mutations in *rpoB* were present in all 12 rifampicin-resistant strains except Vanco + Rif strain 3 (Table 5). Surprisingly, mutation (Ala428Pro) in *rpoB* was also present in MMC + Rif strain 1, which did not display rifampicin resistance. The single amino acid substitution at this position is thus not sufficient to cause resistance.

**Table 5.**
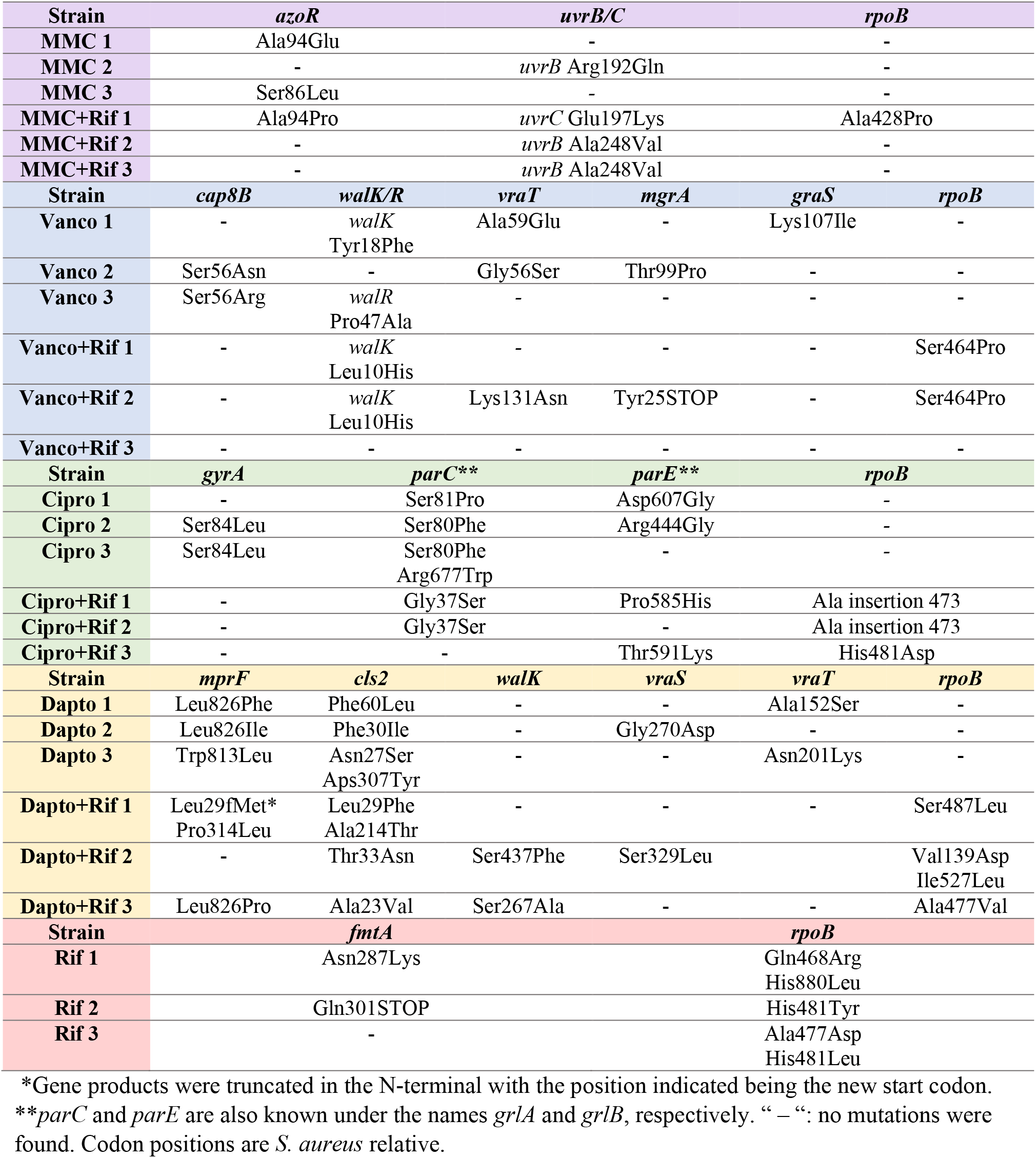
Key mutated genes in evolved strains post adaptive laboratory evolution. Mutations in genes often associated with resistance were searched for by comparing the parent strain pre-evolution with resistant strains post-evolution.

Of all evolved strains, we observed the fewest SNPs in strains evolved under rifampicin monotherapy (figure S1). The mutations were concentrated in *rpoB*, sometimes with more than one mutation in the gene. The only other mutations observed in these strains were in *fmtA*, which encodes the core cell wall stimulon protein, FmtA (34). FmtA is further known to mutate in connection with methicillin resistance in *S. aureus* (35). This finding was surprising, as *fmtA* has not previously been associated with resistance to rifampicin.

As with rifampicin resistance, ciprofloxacin resistance is well characterized and it is often caused by a Ser84Leu mutation in the DNA gyrase subunit A, *gyrA*, resulting in decreased binding affinity of ciprofloxacin to its target (36). We observed this mutation in Cipro strain 2 and Cipro strain 3, which displayed the highest MICs for ciprofloxacin among evolved strains (Table 1). However, these were the only strains with mutations in *gyrA* and ciprofloxacin resistance in the remaining four resistant strains were most likely caused other by mutations in topoisomerase IV subunits *parC*/*grlA* and *parE*/*grlB*, which are also known to be associated with ciprofloxacin resistance (37–39). Although strains with mutations in only *parC/grlA* and *parE/grlB* showed >4 fold increase in MIC we chose as threshold for resistance, the MIC values were much lower than in strains with mutations in *gyrA*. We further observed mutations in *tilS, nrdI, gdpP* and *comEC/rec2*, which have not been connected with ciprofloxacin resistance and which we only observed in single evolved strains (table S1). Since these strains also had mutations in genes with known association to ciprofloxacin resistance, we cannot confirm if they are also linked to ciprofloxacin resistance.

Vancomycin resistance was much more complex than rifampicin and ciprofloxacin resistance. We expected to observe mutations in genes that regulate or contribute to cell wall biosynthesis such, as *vraSRT, walKR, graSR* and *mprF* as demonstrated by others (40–44). The three-component system, VraSRT, is a major regulator of the cell wall stress stimulon, which is initiated after *S. aureus* experiences cell wall stress (45). Likewise, WalKR, is a master regulator of peptidoglycan metabolism and it is required for cell viability (46). The two-component system, GraSR, is mostly known for its implications in vancomycin intermediate resistance (Cui et al., 2009); however, *S. aureus* relies on it during response to cationic antimicrobial peptides (CAMPs) (47). Likewise, MprF is involved in resistance to CAMPs, as it modifies negatively charged phospholipids and introduces positive charges in the membrane (48). We observed mutations in *graS, walK, walR* and *vraT* (*yvqF*), however no particular gene was mutated in all evolved strains, although they reached identical MIC values (Table 1). We further observed mutations in *cap8B* in Vanco strains 2 and 3. *cap8B* is a virulence factor involved in capsule formation (49). Mutations in *cap8B* have not previously been associated with vancomycin resistance in *S. aureus*, but upregulation of *cap8B* has been observed in vancomycin intermediate *S. aureus* (50). Vanco strain 2 and Vanco + Rif strain 2 contained mutations in *mgrA*, a global regulator of virulence and autolysis. *mgrA* has been linked to vancomycin resistance in *S. aureus*, but transposon insertion into *mgrA* only increased vancomycin MIC 2-fold (51) and does not explain the 8-fold increase seen here. Therefore, the increase in MIC must be due to multiple mutations in the same strain rather than a single mutation.

Surprisingly, Vanco + Rif 3 showed no mutations in *cap8B, mgrA* or any of the genes associated with cell wall biosynthesis (Table 5). Instead, we observed a mutation in e.g. *ebh* (Table S1) which is associated with pathogenesis, and mutations in this gene was observed in connection with vancomycin resistance in a single study (40). Furthermore, we observed mutations in *uvrB* and *plsY* (Table S1), which have been shown to be mutated in an *S. aureus* isolate from a patient receiving long term vancomycin treatment (52). We furthermore observed a mutation in *ftsW*, which has not previously been associated with vancomycin resistance, but FtsW and vancomycin both bind lipid II without one preventing binding from the other (53). Combined, the mutations in *ebh, uvrB, plsY* and possibly *ftsW* may be the cause of the vancomycin intermediate phenotype in Vanco + Rif strain 3.

A similarly complex picture emerged from the analysis of daptomycin-resistant evolved strains, which contained a multitude of SNPs. We expected mutations in *mprF, walKR, vraSRT* and *cls2* due to their importance in cell wall biosynthesis and as they have previously been connected to daptomycin non-susceptibility (7, 54, 55). All daptomycin resistant strains contained mutations in *mprF* and *cls2*, except Dapto + Rif strain 2 which did not have mutations in *mprF* (Table 5). Leucine 826 was mutated in 3 of the 6 daptomycin-resistant strains: Dapto strain 1 and 2 (Leu826Phe/Ile), which displayed the highest MIC values (512 μg/mL), and Dapto + Rif strain 3 (Leu826Pro) which had an MIC of (16 μg/mL). Dapto strain 3 had a mutation in a close position (Trp813Leu), and the MIC value was at the >100 μg/mL level (Table 1).

Strains with mutations in the *vraSRT* system were evolved under daptomycin monotherapy and displayed >100 μg/mL MIC values. Dapto strain 2 and Dapto + Rif strain 2 had a mutations in *vraS* while Dapto strains 1 and 3 had mutations in *vraT*. Mutations in *vraT* are known to cause resistance to vancomycin (41) with observations of *vraT* in vancomycin intermediate, dalbavancin non-susceptible *S. aureus* (56). However, there is high occurrence of cross resistance between vancomycin and daptomycin with a linear correlation between the two antibiotics’ MICs (57). Indeed, we observed mutations in the *vraSRT* in both daptomycin- and vancomycin-resistant strains. Like *vraSRT*, we also found mutations in the *walKR* system among both daptomycin-resistant and vancomycin-intermediate strains, as Dapto + Rif strains 2 and 3 had mutations in *walK*.

### Evolution under mitomycin C pressure induces mutations in azoR and uvrB/C

We observed a modest (2-4 fold) increase in the MIC for mitomycin C during evolution under mitomycin exposure as mono- or combination therapy. Although the increase was modest, the change was consistent in all six evolved strains. Mutations that affect bacteria’s susceptibility to mitomycin C have previously been reported in *E. coli*, but only in connection with decreased, not increased, MIC. The gene identified to affect mitomycin C susceptibility was *uvrB* of the nucleotide excision repair (NER) (58, 59). We identified SNPs in the NER-system *uvrABC* in one MMC monotherapy strain and all MMC + Rif strains, with two strains sharing the same mutation in Alanine 248 in *uvrB* (Table 5). Furthermore, three of the six evolved strains contained mutations in the gene of a FMN-dependent NADH-azoreductase, *azoR*. Two strains were mutated in the same position (Alanine 94) and the third strain in a nearby position (Serine 86). As the mutations in Ala 94 in *azoR* and the Ala248Val mutation in *uvrB* both were found in several independently evolved strains, it is highly likely that they are the reason for the increased mitomycin C MIC. However, these genes have not previously been associated with resistance to mitomycin C, and further experiments are needed to verify the connection between these proteins and the potential for developing resistance to mitomycin C.

## Discussion

In this study, we showed that resistance to rifampicin, ciprofloxacin, vancomycin and daptomycin occurs within days or weeks during adaptive laboratory evolution, while mitomycin C was less prone to resistance since the MIC only increased by 2-fold during mitomycin C monotherapy or during mitomycin C + rifampicin combination therapy.

Adaptive evolution under mitomycin C exposure revealed new insights into how bacteria can protect themselves from this drug. The elevated MIC for mitomycin C was most likely linked to mutations in *azoR, uvrB* and *uvrC*. During nucleotide excision repair, UvrABC recognises damaged DNA, cleaves the phosphodiester bond, and subsequently removes 10-15 bp of the damaged DNA which can then be filled out by DNA polymerase I (60). UvrABC has further been implicated in reparation of mono- and interstrand crosslinks following mitomycin C treatment (61). Therefore, it is highly likely that the mutations in *uvrB* and *uvrC* caused the increase in mitomycin C MIC seen here.

Our study identified for the first time the FMN-dependent NADH-azoreductase, *azoR*, as a gene of interest in decreased susceptibility to mitomycin C. There are limited reports on the involvement of *azoR* in antibiotic resistance in *S. aureus*, however, loss-of-function caused resistance against the quinolone JSF-3151 (62). Furthermore, mutations in the AzoR-homologue in *E. coli* conferred resistance to thiol-specific stress from electrophilic quinones and was shown to reduce multiple quinone compounds (63). It has been hypothesized that reducing mitomycin C might modulate its effects, although a direct redox reaction with ascorbic acid did not cause the modulation (64). Therefore, as intracellular reduction of mitomycin C might play a role in protecting the cell against damage, and since the chemical structure of mitomycin C resembles quinones, we hypothesize that AzoR in *S. aureus* may reduce mitomycin C thereby decrease its cytotoxic effects. When electrophilic quinones accumulate on the cell, they can deplete cellular reduced glutathione (GSH). Decreased AzoR activity can therefore also lead to decreased levels of reduced GSH in the cell. Mitomycin C is activated by interacting with GSH or other reactive thiols (65), and AzoR’s effect on the antimicrobial effect on mitomycin C may therefore also work indirectly through its effect on GSH levels. This is the first report of mutations in *azoR* correlating with decreased susceptibility to mitomycin C, and further studies are needed to elucidate the mechanisms of mitomycin C resistance.

We also gained new insights into the evolution of daptomycin resistance, namely that several different mechanisms seem to contribute simultaneously to raise MIC, and that the high MIC also correlate with slower growth rates. Daptomycin is a positively charged lipopeptide drug and the proposed mechanism-of-action is through calcium-dependent insertion into the bacterial membrane with subsequent oligomerization leading to membrane disruption (66). Therefore, one can expect that daptomycin resistance may occur due to changes in the membrane charge or fluidity. Jones et al. demonstrated increased membrane fluidity and net positive surface charge of daptomycin resistant isolates as well as increased translocation of the positively charged lysyl phosphatidylglycerol (LPG) to the membrane in *S. aureus* (67). LPG is synthesized by the bifunctional membrane protein MprF, which further has flippase activity (48). Here, we found that all evolved daptomycin-resistant strains had mutations in the *mprF* gene (Table 5) and it seems most likely that daptomycin resistance occurs due to mutations in *mprF* that cause increased synthesis of LPG, which in turn leads to a more fluid and positively charged membrane that interacts poorly with daptomycin (68).

Evolved strains Dapto strain 1, Dapto strain 2 and Dapto+Rif strain 1 had substantially higher MIC values than the other evolved strains, and they all displayed slower growth rates (Table 1, Figure 3A). Dapto 1 and Dapto 2 had similar mutations in the *mprF* gene with mutations in leucine 826 and Dapto+Rif 1 had a truncated *mprF* with a Pro314Leu mutation. A previous study generated point mutations in *mprF* in a clinical MRSA strain to achieve daptomycin resistance, and these mutations also caused a decrease in growth rate (69). Although there is a connection between mutation of leucine 826 and slow growth for Dapto strain 1 and 2, Dapto + Rif strain 3 had a similar mutation at leucine 826 and did not display slow growth or a high MIC. Therefore, mutation at this amino acid alone is not responsible for high daptomycin resistance or slow growth. The mutations responsible for slowing the growth rate therefore remain to be identified, and our result underlines the complex mechanisms for daptomycin resistance, which relies on several mutations.

The evolution of rifampicin resistance during combination therapy also revealed insights into what drives rifampicin resistance when other antibiotics are used simultaneously. It appears that high rifampicin resistance can evolve even at very low concentrations of rifampicin because a single point mutation can increase MIC values by >100 fold. This was evident in Dapto + Rif 1, which never received more than 8 μg/mL rifampicin during the study, but subsequently had an MIC of 512 μg/mL (Table 1) conferred by a single mutation in *rpoB* (Ser487Leu). To the best of our knowledge, this mutation has not previously been reported in rifampicin resistant *S. aureus*. In contrast, a different mutation in *rpoB* at a nearby location (Ala477Val) in Dapto + Rif strain 3 only caused a moderate (2-fold) increase in MIC.

In general, combination therapy did not prevent the emergence of rifampicin resistance for the antibiotics tested in this study. It was therefore remarkable that strains evolved under mitomycin C and rifampicin combination therapy did not develop resistance to rifampicin. The maximum rifampicin concentration these strains were exposed to was 0.016 μg/mL, and this concentration may therefore be too low to drive rifampicin resistance. However, strains receiving vancomycin and rifampicin combination therapy received maximum 0.063 μg/mL rifampicin, which caused a rifampicin MIC in two of the strains to reach 1024 μg/mL, underlining that a low rifampicin exposure can cause high resistance. Alternatively, spontaneous mutations that confer rifampicin resistance were not selected for in the population when mitomycin C is present. While this is an encouraging result, we must note that the use of rifampicin in the clinic is at higher concentrations, even when used in combination therapy. We can therefore not rule out that rifampicin resistance develops under those conditions, even if rifampicin is combined with mitomycin C. Further research should therefore investigate more deeply the “protective” effect of mitomycin C in relation to rifampicin resistance.

In summary, resistance to rifampicin and the primary antibiotic did develop during combination therapy, however, it was delayed compared to monotherapy. We observed similar genetic mutations for strains receiving combinations as compared with monotherapy e.g. mutations in *azoR* (mitomycin C resistance), *parC/grlA* and *perE/grlB* (ciprofloxacin resistance), *rpoB* (rifampicin resistance) and mutations in genes involved in cell wall biosynthesis (vancomycin and daptomycin resistance). Therefore, combination therapy did not affect the mutation targets and we expect that any strategies to avoid resistance development during monotherapy would further be effective to avoid resistance development during combination therapy.

## Methods

### Strains, growth conditions and antibiotics

*Staphylococcus aureus* ATCC 29213 WT was used to generate evolved antibiotic resistant strains during adaptive laboratory evolution. *S. aureus* was routinely grown overnight in tryptic soy broth (TSB, T8907, Sigma Aldrich) in Erlenmeyer flasks and incubated at 37°C, 180 rpm unless otherwise stated. Antibiotics used for adaptive laboratory evolution and determination of the minimal inhibitory concentration (MIC) were mitomycin C (J63193.MA, Thermo Scientific), vancomycin (Bactocin, MIP Pharma GmbH), ciprofloxacin (17850, Sigma Aldrich), daptomycin (Cubicin, Merck Sharp & Dohme) and rifampicin (Rifadin, Sanofi S.r.I.).

### Minimum inhibitory concentration determination

MICs were determined using broth dilution in TSB, but otherwise following the ISO standard (70). Briefly, *S. aureus* was diluted to a turbidity of 0.05 and inoculated in 2-fold antibiotic dilution series in TSB in 96-well plates, yielding a bacterial concentration of 5 × 10^5^ CFU/mL. Plates were incubated overnight at 37 °C, 50 rpm and optical density at 600 nm (OD_600_) was measured to detect growth. MIC was determined as the lowest antibiotic concentration with OD_600_ < 20% of the growth control. 50 μg/mL Ca^2+^ was added in samples with daptomycin.

### Adaptive laboratory evolution assay

Three independent colonies (parent 1, 2 and 3) were inoculated in TSB and incubated overnight. The three samples were diluted to a turbidity of 0.05 and inoculated in antibiotic dilution series in 96-well plates. The antibiotic treatments included mitomycin C, vancomycin, ciprofloxacin, daptomycin, rifampicin, mitomycin C + rifampicin, vancomycin + rifampicin, ciprofloxacin + rifampicin and daptomycin + rifampicin. 96-well plates were incubated overnight at 37 °C and 50 rpm and subsequently the MIC was determined by reading the plates in a plate reader at 600 nm. Bacteria from the dilution step immediately below MIC were transferred to a fresh dilution series of antibiotic treatments and incubated overnight. The range of antibiotic concentrations used was from 0.25 to 2-fold the MIC value from the previous day. This cycle was repeated 42 times. As the MIC increased, bacteria surviving higher antibiotic concentrations were transferred to the next cycle with increased antibiotic concentrations, thereby adapting the assay to the development of resistance. After 42 cycles, evolved strains were grown overnight on TSB agar without antibiotics before storage at -80 °C in 25% glycerol. Evolved strains were named after the treatment they received during adaptive laboratory evolution i.e. MMC, Vanco, Cipro, Dapto, Rif, MMC + Rif, Vanco + Rif, Cipro + Rif and Dapto + Rif. In combination treatments, the relative concentration of rifampicin to the primary antibiotic was held constant throughout the adaptive evolution. For each, three independently evolved strains were generated, yielding 27 uniquely evolved strains.

### Growth curves

Overnight cultures of parent and evolved strains were diluted 1000-fold in TSB and incubated in duplicate in a 96-well plate in a shaking plate reader at 37 °C, where plates were shaken for 10 sec at 200 rpm immediately before each measurement. OD_600_ was measured every 30 min for 24 h.

### Biofilm formation

Overnight cultures of parent and evolved strains were diluted to OD_600_ = 1 in TSB and added to 96 well plates with peg-lids pre-conditioned with TSB. After 30 min attachment, peg-lids were transferred to fresh TSB and 96-well plates were incubated at 37 °C, 50 rpm for three days with exchange of media every 24 h. Subsequently, peg-lids were washed twice by transferring to 1 × M9 salts (M6030, Sigma Aldrich) for 30 s, twice. The peg lids were sonicated in M9 salts for 10 min to detach biofilms, and CFU enumeration was subsequently performed on the sonicate.

### Genome sequencing, assembly and bioinformatics analyses

DNA was extracted from overnight cultures using DNeasy UltraClean Microbial Kit following the manufacturer’s protocol (QIAGEN, 12224-50) and prepared for sequencing using Nextera XT DNA Library prep kit (Illumina, FC-131-1024) to tagment the DNA with adapter sequences. The DNA was amplified and adapter and index sequences were added through PCR. AMPure XP magnetic beads were used for DNA purification, removal of primers and short fragments. Finally, the sequences were sequenced on an Illumina MiSeq Next Generation sequencer.

Sequenced genomes were quality controlled using FastQC (71) and trimmed based on per base sequence content using trimmomatic (72). Genomes were assembled and mapped from a reference genome (NCBI: *Staphylococcus aureus* ATCC 29213, assembly GCA_001267715.2) using BactSNP (73). Assemblies were quality controlled using Quast (74) and SNP annotation was performed using SnpEff (75). In order to validate the located SNPs, genomes were further annotated with Prokka (76) and searches for specific mutations were conducted using BLASTp (77).

### Statistical analyses

Ordinary one-way ANOVA was used for bar graphs with a post-hoc uncorrected Fisher’s test. Absolute values of the minimum inhibitory concentration is shown for a minimum of three replicates. GraphPad Prism was used for all statistical analyses (v. 9.5.1 (733) for Windows, GraphPad Software, San Diego, California USA, www.graphpad.com).

## Acknowledgements

We acknowledge Maria Braad Lund and the technical staff at the Department of Biology – Microbiology at Aarhus University for their expertise and assistance in DNA extraction and genome sequencing. We acknowledge Ian Marshall for his assistance during the bioinformatics analysis of SNP calling and annotation. This work was supported by the Novo Nordisk Foundation (grant no. NNF19OC0058357). The funders had no role in study design, data collection and interpretation, or the decision to submit the work for publication.

